# Proprioceptive afferents differentially contribute to effortful perception of object heaviness and mass distribution

**DOI:** 10.1101/2020.07.18.210401

**Authors:** Madhur Mangalam, Nisarg Desai, Tarkeshwar Singh

**Affiliations:** Department of Physical Therapy, Movement and Rehabilitation Science, Northeastern University, Boston, MA, USA; Department of Anthropology, University of Minnesota, Minneapolis, MN, USA; Department of Kinesiology, University of Georgia, Athens, GA, USA

**Keywords:** inertia perception, mass perception, proprioception, kinesthesia, dynamic touch, psychophysics

## Abstract

When humans handle a tool, such as a tennis racket or hammer, for the first time, they often wield it to determine its inertial properties, however, the mechanisms that contribute to perception of inertial properties are not fully understood. The goal of the present study was to investigate how proprioceptive afferents contribute to effortful perception of heaviness and mass distribution of a manually wielded object in the absence of vision. Blindfolded participants manually wielded a set of specially-designed experimental objects of different mass and mass distribution about the wrist at different wrist angles and wrist angular kinematics. By independently manipulating these variables, we aimed to elicit different levels of tonic and rhythmic activity in the muscle spindles of the wrist flexors and extensors and relate them to reported perceptual judgments of heaviness and length. Perception of heaviness and length were predominantly dependent on an object’s static moment and the moment of inertia, respectively. Manipulations of wrist angle and wrist angular kinematics affected perceptual judgments of heaviness and length in relatively opposite ways. As for wrist angle, ulnar deviation consistently resulted in an object being perceived heavier but shorter. Compared to static holding, wielding the object resulted in it being perceived heavier but wielding did not affect perceived length. These results suggest that proprioceptive afferents differentially contribute to effortful perception of object heaviness and mass distribution.

## Introduction

Our performance in everyday activities involving handheld objects relies on perception of properties of those objects, such as heaviness, length, and shape, often explicitly through proprioceptive feedback. For instance, the heaviness of a hammer, the length of a stick, and the mass distribution of a tennis racket, all can be perceived by wielding the respective object about the wrist. Movement generates proprioceptive and kinesthetic afferent feedback that contributes to perception of object properties. The neurophysiological basis of how afferent feedback contributes to stable perception of object properties remains unknown. The goal of this study was to investigate how kinesthetic feedback (of limb position and movement) from muscle spindles and force feedback from Golgi tendon organs contribute to perception of two distinct properties of handheld objects: heaviness and length.

Until recently, the conventional understanding was that perception of heaviness is solely based on an efference copy of motor commands sent by the motor cortex to the somatosensory areas. This “central effort” hypothesis was supported by the findings that perceived heaviness increases after muscle fatigue despite no accompanying changes in afferent activity (Gandevia and McCloskey 1977a, b; Aniss et al. 1988; Proske and Allen 2019), and blocking afferent feedback by anesthesia does not affect perception of heaviness (Gandevia and McCloskey 1977a, c; Proske and Gandevia 2012; Proske and Allen 2019). However, this hypothesis has been questioned, as it fails to reconcile recent findings on the contribution of afferent feedback to perception of heaviness (Luu et al. 2011; Brooks et al. 2013; Proske and Allen 2019). A model for heaviness perception based on fusimotor reafference has been proposed to reconcile these findings (Luu et al. 2011).

The putative role of feedback from spindles and tendon organs is much essential to perception of object properties specified by mass distribution than for heaviness, which can be perceived with reasonable accuracy even when all peripheral feedback is blocked by anesthesia. For instance, one can judge which one of the rackets is heavier by supporting them in hand in the absence of any movement but we need to wield them to judge if a given racket is head-light or head-heavy. A body of research on effortful perception has been founded on the hypothesis that the definite scaling of the non-visual perceptions of spatial dimensions (e.g., length, width, and shape) and other properties (e.g., orientation in hand) of a manually wielded object has its basis in the moments of the object’s mass distribution (Carello and Turvey 2000; Turvey and Carello 2011). It seems that as opposed to perception of heaviness, perception of object properties related to mass distribution relies more on feedback from spindles than from tendon organs. It has been found, for example, that people can perceive the length of an occluded object by wielding in hand with reasonable accuracy even when that object is immersed in water (Pagano and Donahue 1999; Pagano and Cabe 2003; Mangalam et al. 2017, 2018), and the force of buoyancy reduces the force required to wield that object, the central effort, and the peripheral feedback from tendon organs. These studies suggest that feedback from spindles and tendon organs may differentially contribute to the effortful perception of object heaviness and length. However, how the afferent feedback from spindles and tendon organs interact to produce the perception of heaviness and length remains elusive.

The goal of the present study was to parse out the contributions of feedback from spindles and tendon organs in the effortful perception of heaviness and length of manually wielded objects in the absence of vision. A significant technical problem limiting the scope of neurophysiological investigations of peripheral afferent feedback in healthy humans is that the spindle and tendon organ activity cannot be measured directly without using invasive techniques such as microneurography. Studies have typically relied on vibrations of different frequencies to selectively modulate the feedback from spindles and tendon organs (Fallon and Macefield 2007; Luu et al. 2011; Brooks et al. 2013). An alternative approach is to examine the effects of the manipulations of joint angle positions and kinematics—which modulate the feedback from both spindles and tendon organs, respectively (Proske and Gandevia 2012), on perception. We asked blindfolded participants to manually wield specially-designed experimental objects of different mass and mass distribution about the wrist at different wrist angles and kinematics and report their perceptual judgments of heaviness and length. Our specific hypothesis in this study was that changes in static and dynamic wrist joint kinematics during object wielding would affect perception of object heaviness and length.

## Methods

### Participants

Seven adult men and five adult women (*M*±1*SD* age = 25±0.8 years, right-handed) voluntarily participated in the present experiment after providing written consent approved by the Institutional Review Board (IRB) at the University of Georgia (Athens, GA).

### Experimental model

Consider a simplified two-dimensional task of wielding a weightless rod having a point mass *m* attached at a distance *d* to the wrist joint, such that:

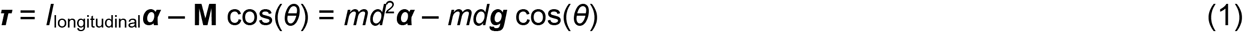

where ***τ*** is the muscular torque, *I*_longitudinal_ reflects the resistance of the object to rotational movement about the wrist along the longitudinal axis, ***α*** is the angular acceleration of the wielded object, ***g*** is the gravitational acceleration, *θ* is the angle of the object relative to the horizontal plane, and *d* is the distance of the point mass *m* to the wrist. The right-hand side of Eq. 1 includes the moment of inertia, *I*_longitudinal_ = *md*^2^ and the static moment **M** (= *md**g***).

Although both the static moment and the moment of inertia describe mass distribution—both depend on mass (*m*) and position of that mass (*d*)—the two parameters have distinct implications for perception. The moment of inertia can influence perception only through angular acceleration, whereas the static moment can influence perception at rest as well. Additionally, the static moment shows a linear dependence on *d*, whereas the moment of inertia shows a quadratic dependence on *d*. Thus, the contribution of one of the two parameters can be controlled by holding the other parameter constant. *I*_longitudinal_ can be held constant while varying **M** by increasing *m* fourfold and halving *d*. **M** can be held constant while varying *I*_longitudinal_ by doubling *m* and halving *d*. Accordingly, to investigate which object parameter provides the basis for perception of heaviness and length, we designed six experimental objects that systematically differed in mass, the static moment, and the moment of inertia.

### Experimental objects

Each participant wielded six experimental objects, each object consisting of a dowel (oak, hollow aluminum, or solid aluminum; diameter = 1.2 cm, length = 75.0 cm) weighted by 4 or 12 stacked steel rings (inner diameter = 1.4 cm, outer diameter = 3.4 cm, thickness = 0.2 cm, mass = 14 g) attached to the dowel at 20.0 or 60.0 cm, respectively (Table 1 and Fig. 1A). The dowels were weighted such that the resulting six objects systematically differed in mass, *m* (Object 1 < Object 2, Object 3 < Object 4, Object 5 < Object 6), the static moment, **M** (Object 1 = Object 2 = **M**_S_ < Object 3 = Object 4 = **M**_M_ < Object 5 = Object 6 = **M**_L_), and the moment of inertia, *I*_longitudinal_ (Object 1 > Object 2, Object 3 > Object 4, Object 5 > Object 6). A cotton tape of negligible mass was enfolded on each dowel to prevent the cutaneous perception of its composition (i.e., oak versus aluminum).

**Table 1.**
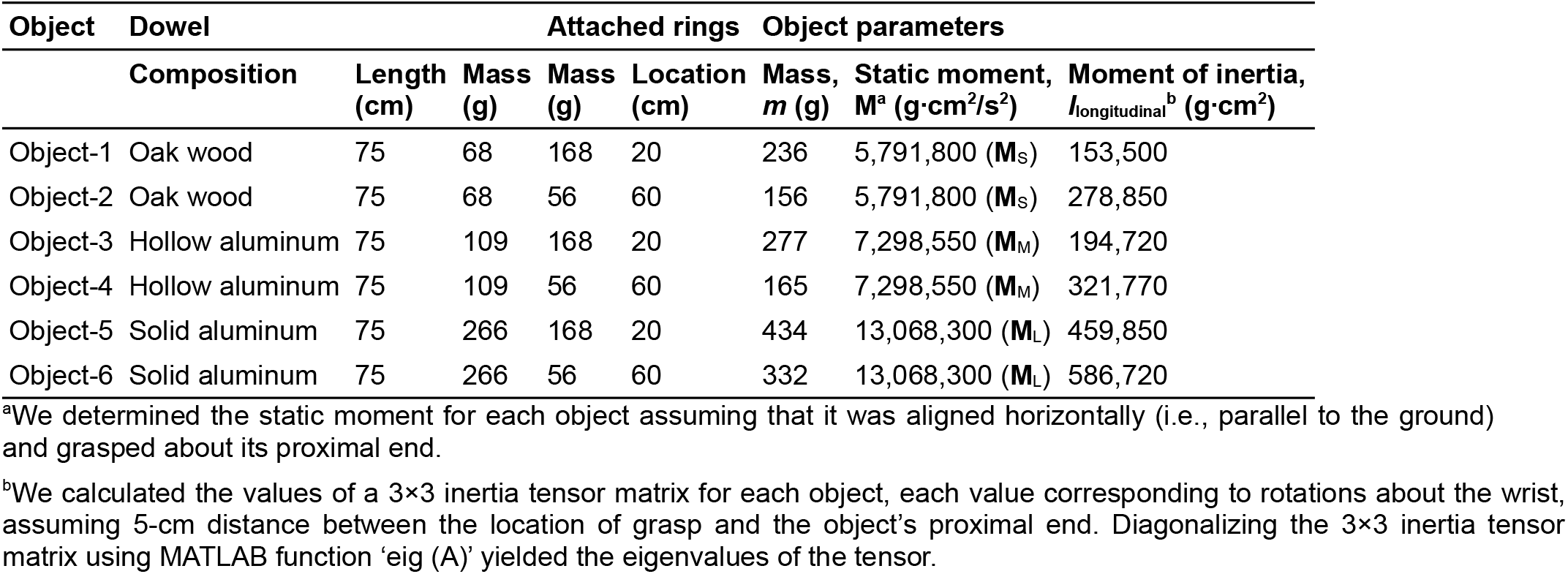
Experimental objects

**Fig 1.**
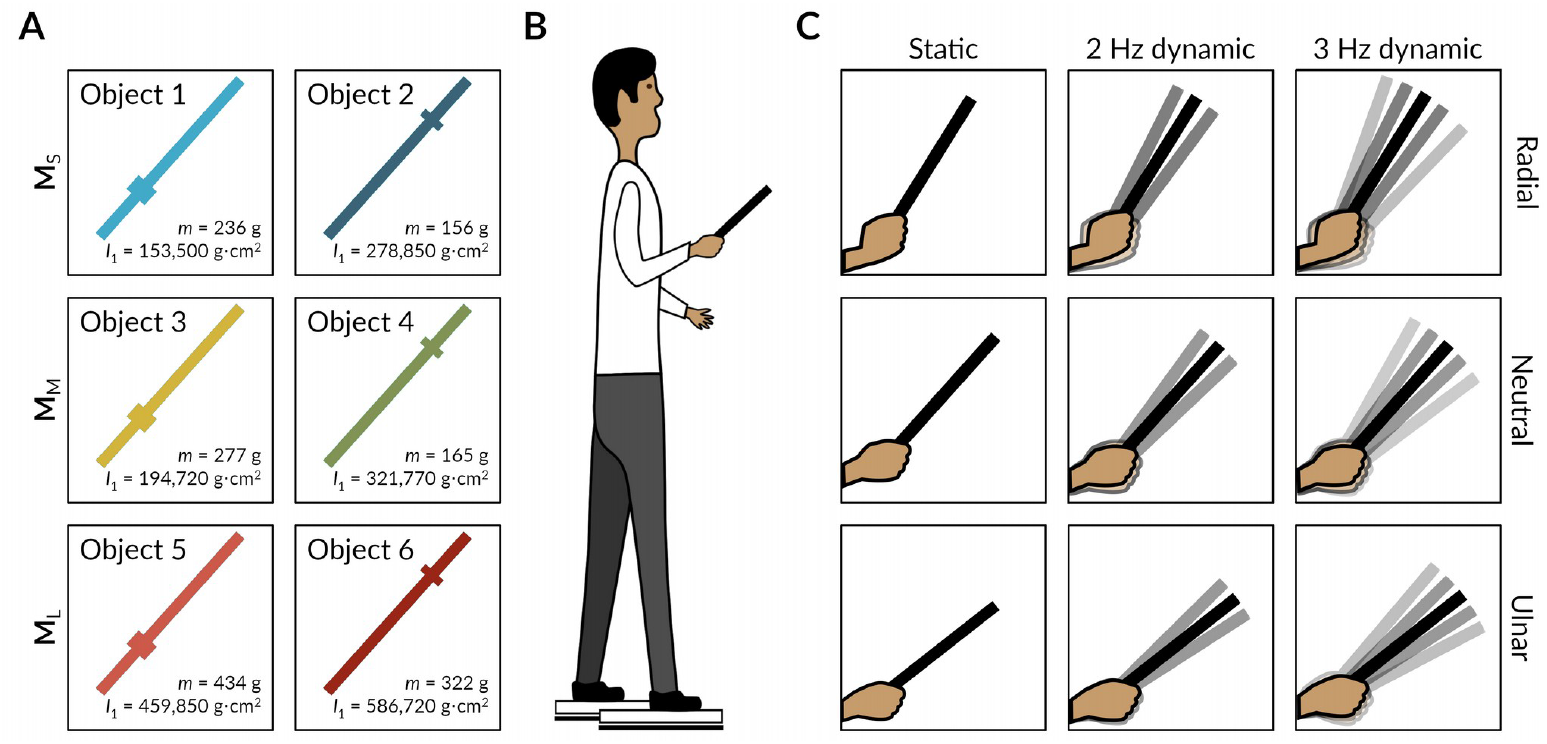
Schematic illustration of the experimental objects, setup, and exploratory conditions. (**A**) Each participant wielded six different weighted dowels that systematically differed in mass, *m* (Object-1 < Object-2, Object-3 < Object-4, Object-5 < Object-6), the static moment, **M** (**M**_S_ < **M**_M_ < **M**_L_), and the moment of inertia, *I*_longitudinal_ (Object-1 > Object-2, Object-3 > Object-4, Object-5 > Object-6). (**B**) Each participant wielded each object for 5 s and reported their judgments of the heaviness (with respect to a reference object) and the length of that object. (**C**) The participant was instructed to constrain his/her wrist movement about 10° radial deviation (top panels), a neutral position (middle panels), or about 10° ulnar deviation (bottom panels). In a static condition, the participant lifted and held each object static (left panels), and in the two dynamic conditions, the participant lifted and wielded each object synchronously with metronome beats at 2 Hz or 3 Hz (center and right panels).

### Experimental task, procedure, and instructions to the participants

Feedback from the spindles and tendon organs was differentially manipulated by asking participants to wield objects at different wrist angles and kinematics. Each participant was asked to wield each object at three different wrist angles: (1) 10° radial deviation (Fig. 1C, top panels), (2) neutral position (Fig. 1C, middle panels), and (3) 10° ulnar deviation (Fig. 1C, bottom panels). We expected that the ulnar and radial deviations of the wrist would increase the [baseline] spindle and tendon organ activity in the antagonist muscles: the radial and ulnar muscles of the hand, respectively. Additionally, at each wrist angle, each participant was asked to wield each object about the wrist at different angular speeds. In a static condition, each participant was asked to lift and hold each object (Fig. 1C, left panels). In the two dynamic conditions, each participant was asked to lift and wield each object synchronously with metronome beats at 2 Hz or 3 Hz (Fig. 1C, center and right panels). At any given wrist angle, wielding an object at different speeds would modulate the reafference from the spindles in addition to modulating the tendon organ activity—faster movement will result in increased spindle reafference. Each participant was instructed to wield the object at small amplitude so as to maintain the wrist angle at the ulnar, neutral, and radial positions through the entire trial.

Each participant stood on a designated location and assumed a given wrist angle comfortably. A custom setup consisting of two tripods was used to support and align each experimental object relative to the participant’s wrist at the ulnar, neutral, and radial positions (this setup is not shown in Fig. 1). Changing the heights of these two tripods—one lower and the other higher relative to the participant’s hand—allowed us to present an object, so the participant readily held that object at the ulnar, neutral, and radial positions of the wrist upon grasping it. At the beginning and after every six trials, the participant wielded a reference object (an unweighted hollow aluminum dowel, diameter = 1.2 cm, length = 75.0 cm, mass = 109 g) and assigned it a heaviness value of 100. He/she assigned heaviness values proportionally higher than 100 to an object perceived heavier than the reference object (e.g., 200 to an object perceived twice as heavy), and heaviness values proportionally less than 100 to that perceived lighter than the reference object (e.g., 50 to an object perceived half as heavy). In each trial, following a ‘lift’ signal, the participant lifted the object and held it static or wielded it synchronously with metronome beats at 2 Hz or 3 Hz. After 5 s and following a ‘stop’ signal, the participant kept the object back and reported perceived heaviness relative to the reference object and perceived length by changing the position of a marker by pulling a string on a string-pulley assembly. We instructed the participant to minimized the movement amplitude in the 2 Hz and 3 Hz dynamic conditions. The 5 s duration was chosen to minimize memory-based comparisons from previous trials.

Each participant was tested individually in a 90–105-min session during which he/she completed a total of 108 trials: 3 Wrist angles × 3 Wrist angular kinematics × 6 Objects × 2 Trials. A crossed, pseudo-randomized block design was used, the factors of Wrist angle (Radial, Neutral, and Ulnar) crossed with the factors of Wrist angular kinematics (Static, 2 Hz dynamic, and 3 Hz dynamic). The order of the 12 trials (6 Objects × 2 Trials/Object) was pseudo-randomized for each block.

### Statistical analysis

To investigate which object parameters best explained variation in perceived heaviness and length, we followed an information-theoretic approach. This approach uses the Akaike Information Criterion (AIC; or quasi-AIC (QAICc) for over-dispersed data) to choose a set of plausible models from a given set of *a priori* candidate models (Burnham and Anderson 2002). According to this approach, the Akaike information criterion (AIC) serves as an estimator of out-of-sample prediction error and thereby the relative quality of statistical models for a given set of data. AIC estimates the quality of each model relative to each of the other models. Specifically, a smaller AIC value reflects better performance/complexity trade-off. Thus, AIC provides a means for selecting the model with the best performance/complexity trade-off. We considered eight candidate models, including the null model and all the different combinations of the given object parameters: mass, the static moment, and the logarithm of moment of inertia. We scaled the object parameters to a mean of 0 and a standard deviation of 1 to aid model fitting and avoid the effects of scale. We performed this analysis for heaviness and length separately and controlled for participant identity in each model using linear mixed-effects models (LMEs).

To examine the effects of wrist angle, wrist angular kinematics, and object on perception, we submitted the values of perceived heaviness and perceived length to aligned rank transformed (ART) ANOVA, using the *‘ARTool’ R*-package in *R*. ART ANOVA is a nonparametric approach that accommodates multiple independent variables, interactions, and repeated measures. Significant main and interaction effects were followed by post-hoc comparisons with the *p*-values corrected for multiple comparisons using Tukey’s method for pairwise contrasts and Holm method for interaction contrasts. Each test statistic was considered significant at the two-tailed alpha level of 0.05.

Because our data did not fit any exponential family distribution, it required a non-parametric approach like the ART ANOVA (using ranked data) for inference. However, we still report Cohen’s *d*-like effect sizes—approach as taken in Rouder et al. (2012)—on the actual data (not ranked data) from the linear mixed-effects model (LME) using the ‘nlme’ R-package in R. Although the LMEs will be less reliable given the data distribution, they can still be used to make sense of the effect sizes.

## Results

### Distinct object parameters specified perceptual judgments of heaviness and length

To investigate which object parameters best explained variation in perceived heaviness and length, we followed an information-theoretic approach to model selection. Of the eight models we considered (Table 2), four of the five models with non-zero probability included the static moment; the fifth model included mass and the moment of inertia but not the static moment. The model with the best performance-to-complexity ratio (i.e., smallest QAICc value) included only the static moment, and the support for this model was 1.46 times stronger than the model also including mass (evidence ratio = *W*_i_/*W*_j_ = 0.35/0.24 = 1.46; Table 2 and Fig. 2A, top panel), 1.84 times stronger than the model also including the moment of inertia (*W*_i_/*W*_j_ = 0.35/0.19 = 1.84), and 3.89 times stronger than the model including all object parameters (*W*_i_/*W*_j_ = 0.35/0.09 = 3.89). The second-best model also included object mass, which is consistent with the finding that for each value of the static moment, the object with a greater mass was perceived to be heavier (Tables 3 & 4, and Fig. 2A, bottom panel). Although all four models that also included mass and/or the moment of inertia showed closer fits, as reflected by the smaller log-likelihood values, the improvement in fit was accompanied by an increase in the complexity of the model, ultimately reducing the performance-to-complexity ratio, as reflected by the larger QAICc values. In other words, including mass and/or the moment of inertia in a model likely overfitted the data than it increased its predictive power. Considering model-averaged parameter estimates (Burnham and Anderson 2002), an increase in the static moment resulted in an increase in perceived heaviness; for the other two object parameters, the 95% confidence interval set included zero (Table 2).

**Fig 2.**
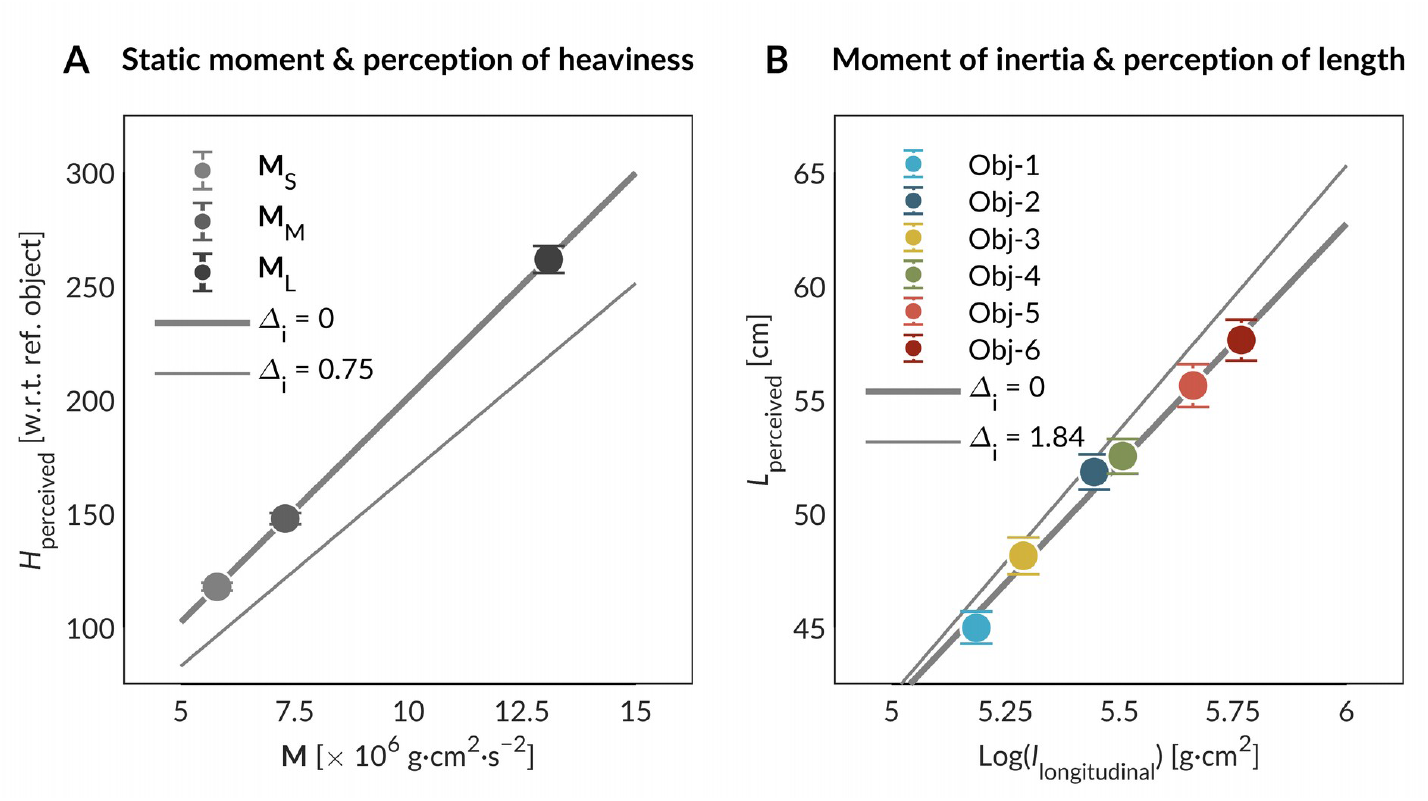
Perception of heaviness and length via effortful touch was based on distinct object parameters. (**A**) Perceived heaviness increased as a function of the static moment. (**B**) Perceived length increased as a function of the moment of inertia. The thicker and thinner lines in the top panels indicate the best and the second-best model fits, respectively. Error bars indicate 1SEM. Mass: Object-1 > Object-2, Object-3 > Object-4, Object-5 > Object-6; Static moment: **M**_S_ < **M**_M_ < **M**_L_; Moment of inertia: Object-1 < Object-2, Object-3 < Object-4, Object-5 < Object-6.

**Table 2.**
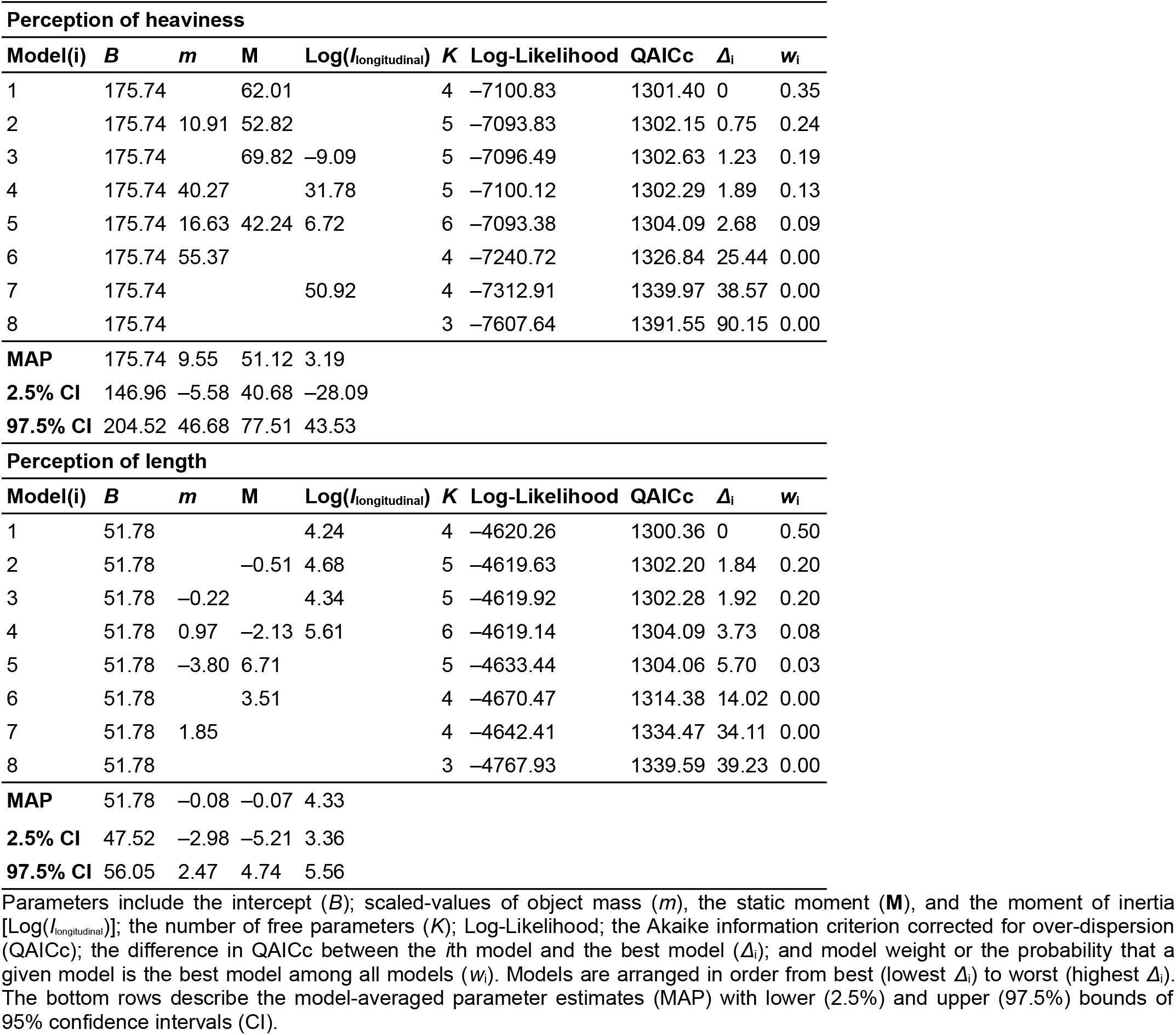
Summary of model selection

| Perception of heaviness |  |  |  |  |  |  |  |  |  |
| --- | --- | --- | --- | --- | --- | --- | --- | --- | --- |
| Model(i) | *B* | *m* | *M* | Log(*l*<sub>longitudinal</sub>) | *K* | Log-Likelihood | QAICc | $\Delta_i$ | *w<sub>i</sub>* |
| 1 | 175.74 |  | 62.01 |  | 4 | −7100.83 | 1301.40 | 0 | 0.35 |
| 2 | 175.74 | 10.91 | 52.82 |  | 5 | −7093.83 | 1302.15 | 0.75 | 0.24 |
| 3 | 175.74 |  | 69.82 | −9.09 | 5 | −7096.49 | 1302.63 | 1.23 | 0.19 |
| 4 | 175.74 | 40.27 |  | 31.78 | 5 | −7100.12 | 1302.29 | 1.89 | 0.13 |
| 5 | 175.74 | 16.63 | 42.24 | 6.72 | 6 | −7093.38 | 1304.09 | 2.68 | 0.09 |
| 6 | 175.74 | 55.37 |  |  | 4 | −7240.72 | 1326.84 | 25.44 | 0.00 |
| 7 | 175.74 |  |  | 50.92 | 4 | −7312.91 | 1339.97 | 38.57 | 0.00 |
| 8 | 175.74 |  |  |  | 3 | −7607.64 | 1391.55 | 90.15 | 0.00 |
| MAP | 175.74 | 9.55 | 51.12 | 3.19 |  |  |  |  |  |
| 2.5% CI | 146.96 | −5.58 | 40.68 | −28.09 |  |  |  |  |  |
| 97.5% CI | 204.52 | 46.68 | 77.51 | 43.53 |  |  |  |  |  |
| Perception of length |  |  |  |  |  |  |  |  |  |
| Model(i) | *B* | *m* | *M* | Log(*l*<sub>longitudinal</sub>) | *K* | Log-Likelihood | QAICc | $\Delta_i$ | *w<sub>i</sub>* |
| 1 | 51.78 |  |  | 4.24 | 4 | −4620.26 | 1300.36 | 0 | 0.50 |
| 2 | 51.78 |  | −0.51 | 4.68 | 5 | −4619.63 | 1302.20 | 1.84 | 0.20 |
| 3 | 51.78 | −0.22 |  | 4.34 | 5 | −4619.92 | 1302.28 | 1.92 | 0.20 |
| 4 | 51.78 | 0.97 | −2.13 | 5.61 | 6 | −4619.14 | 1304.09 | 3.73 | 0.08 |
| 5 | 51.78 | −3.80 | 6.71 |  | 5 | −4633.44 | 1304.06 | 5.70 | 0.03 |
| 6 | 51.78 |  | 3.51 |  | 4 | −4670.47 | 1314.38 | 14.02 | 0.00 |
| 7 | 51.78 | 1.85 |  |  | 4 | −4642.41 | 1334.47 | 34.11 | 0.00 |
| 8 | 51.78 |  |  |  | 3 | −4767.93 | 1339.59 | 39.23 | 0.00 |
| MAP | 51.78 | −0.08 | −0.07 | 4.33 |  |  |  |  |  |

|  |  |  |  |  |
| --- | --- | --- | --- | --- |
| **2.5% CI** | 47.52 | -2.98 | -5.21 | 3.36 |
| **97.5% CI** | 56.05 | 2.47 | 4.74 | 5.56 |

Parameters include the intercept ( $B$ ); scaled-values of object mass ( $m$ ), the static moment ( $\mathbf{M}$ ), and the moment of inertia [ $\text{Log}(I_{\text{longitudinal}})$ ]; the number of free parameters ( $K$ ); Log-Likelihood; the Akaike information criterion corrected for over-dispersion (QAICc); the difference in QAICc between the $i$ th model and the best model ( $\Delta_i$ ); and model weight or the probability that a given model is the best model among all models ( $w_i$ ). Models are arranged in order from best (lowest $\Delta_i$ ) to worst (highest $\Delta_i$ ). The bottom rows describe the model-averaged parameter estimates (MAP) with lower (2.5%) and upper (97.5%) bounds of 95% confidence intervals (CI).

**Table 3.** Coefficients of ART ANOVA examining the influence of wrist angle, wrist angular kinematics, and object on perceived heaviness and length

| Effect | Perceived heaviness |  |  | Perceived length |  |  |
| --- | --- | --- | --- | --- | --- | --- |
|  | *F* | Df, Df.res | *P*<sup>a</sup> | *F* | Df, Df.res | *P*<sup>a</sup> |
| Wrist angle | 23.56 | 2, 1231 | **< 0.001** | 13.37 | 2, 1231 | **< 0.001** |
| Wrist angular kinematics | 8.05 | 2, 1231 | **< 0.001** | 1.36 | 2, 1231 | 0.257 |
| Object | 456.87 | 5, 1231 | **< 0.001** | 66.21 | 5, 1231 | **< 0.001** |
| Wrist angle × Wrist angular kinematics | 4.51 | 4, 1231 | **0.001** | 5.66 | 4, 1231 | **0.001** |
| Wrist angle × Object | 1.55 | 10, 1231 | 0.118 | 0.75 | 10, 1231 | 0.676 |
| Wrist angular kinematics × Object | 1.13 | 10, 1231 | 0.339 | 0.85 | 10, 1231 | 0.583 |
| Wrist angle × Wrist angular kinematics × Object | 0.66 | 20, 1231 | 0.870 | 0.99 | 20, 1231 | 0.476 |

<sup>a</sup>Boldfaced values indicate statistical significance at the two-tailed alpha level of 0.05.

**Table 4.** Pairwise contrasts following ART ANOVAs in Table 3.

|  |  | Perceived heaviness |  |  |  | Perceived length |  |  |  |
| --- | --- | --- | --- | --- | --- | --- | --- | --- | --- |
| Effect | Contrast | Estimate ($\pm 1SE$) | Df | t-ratio | P<sup>a</sup> | Estimate ($\pm 1SE$) | Df | t-ratio | P<sup>a</sup> |
| Wrist angle |  |  |  |  |  |  |  |  |  |
|  | Neutral – Ulnar | −90.8 (18.6) | 1231 | −4.88 | < **0.001** | 67.5 (19) | 1231 | 3.55 | **0.001** |
|  | Radial – Ulnar | −123.3 (18.6) | 1231 | −6.62 | < **0.001** | 97.5 (19) | 1231 | 5.03 | < **0.001** |
|  | Radial – Neutral | −32.4 (18.6) | 1231 | −1.74 | 0.190 | 28.2 (19) | 1231 | 1.49 | 0.299 |
| Wrist angular kinematics |  |  |  |  |  |  |  |  |  |
|  | 2 Hz dynamic – Static | 73.9 (18.7) | 1231 | 3.96 | < **0.001** | 22 (19.1) | 1231 | 1.15 | 0.482 |
|  | 3 Hz dynamic – Static | 47.4 (18.7) | 1231 | 2.54 | **0.030** | 8.48 (19.1) | 1231 | 0.45 | 0.897 |
|  | 3 Hz dynamic – 2 Hz dynamic | −26.5 (18.7) | 1231 | −1.42 | 0.332 | 39.48 (19.1) | 1231 | 1.60 | 0.247 |
| Effect | Contrast | Value | Df | $\chi^2$ | P<sup>a</sup> | Value | Df | $\chi^2$ | P<sup>a</sup> |
| Wrist angle × Wrist angular kinematics |  |  |  |  |  |  |  |  |  |
|  | Neutral – Ulnar × Static – 2 Hz dynamic | 32.90 | 1 | 0.52 | 1.000 | 116.64 | 1 | 6.33 | 0.059 |
|  | Neutral – Ulnar × Static – 3 Hz dynamic | −118.23 | 1 | 6.71 | 0.058 | −59.79 | 1 | 1.66 | 0.592 |
|  | Neutral – Ulnar × 2 Hz dynamic – 3 Hz dynamic | −151.13 | 1 | 10.96 | **0.007** | −176.42 | 1 | 14.48 | **0.001** |
|  | Radial – Ulnar × Static – 2 Hz dynamic | −22.94 | 1 | 0.25 | 1.000 | −17.08 | 1 | 0.14 | 0.712 |
|  | Radial – Ulnar × Static – 3 Hz dynamic | −154.85 | 1 | 11.51 | **0.006** | −138.49 | 1 | 8.92 | **0.023** |
|  | Radial – Ulnar × 2 Hz dynamic – 3 Hz dynamic | −131.90 | 1 | 8.35 | **0.027** | −121.41 | 1 | 6.86 | 0.053 |
|  | Radial – Neutral × Static – 2 Hz dynamic | −55.84 | 1 | 1.49 | 1.000 | −133.72 | 1 | 8.32 | **0.028** |
|  | Radial – Neutral × Static – 3 Hz dynamic | −36.62 | 1 | 0.64 | 1.000 | −78.70 | 1 | 2.88 | 0.359 |
|  | Radial – Neutral × 2 Hz dynamic – 3 Hz dynamic | 19.22 | 1 | 0.18 | 1.000 | 55.01 | 1 | 1.41 | 0.592 |

<sup>a</sup>Boldfaced values indicate statistical significance at the two-tailed alpha level of 0.05.

An identical model comparison yielded very distinct effects of the three object parameters on judgments of length. Of the eight models we considered (Table 2), four of the five models with non-zero probability included the moment of inertia; the fifth model included mass and the static moment but not the moment of inertia. The model with the best performance-to-complexity ratio (i.e., smallest QAICc value) included only the moment of inertia, and the support for this model was 2.50 times stronger than the model also including mass (evidence ratio = *W*_i_/*W*_j_ = 0.50/0.20 = 2.50; Table 2 and Fig. 2B, top panel), 2.50 times stronger than the model also including the static moment (*W*_i_/*W*_j_ = 0.50/0.20 = 2.50), and 6.25 times stronger than the model including all three object parameters (*W*_i_/*W*_j_ = 0.50/0.08 = 6.25). In contrast to perception of heaviness, for each value of the static moment, the object with a greater moment of inertia—and not mass—was perceived to be longer (Tables 3 & 4, and Fig. 2B, bottom panel). Although all four models that also included mass and the static moment showed closer fits, as reflected by the smaller log-likelihood values, the improvement in fit was accompanied by an increase in the complexity of the model, ultimately reducing the performance-to-complexity ratio, as reflected by the larger QAICc values. In other words, including mass and/or the static moment in a model likely overfitted the data as opposed to increasing its predictive power. Considering model-averaged parameter estimates (Burnham and Anderson, 2002), an increase in the moment of inertia resulted in an increase in perceived length; for the other two object parameters, the 95% confidence interval set included zero (Table 2).

These results suggest that the effortful perception of heaviness and length of an occluded wielded object was based on distinct object parameters—perceived heaviness of an object was a function of its static moment, and perceived length was a function of its moment of inertia. (Note that the actual length of all six objects was 75 cm.) These results reflect the everyday experience with perception of object properties: given that the static moment of an object about the point of rotation depends on the force of gravity, an object is perceived lighter when immersed in water than when wielded in the air. In contrast, given that the moment of inertia of an object does not depend on the external forces acting on that object, perceived length of an object is fairly consistent whether that object is wielded in water or the air, as several studies have shown previously (Pagano and Donahue 1999; Pagano and Cabe 2003; Mangalam et al. 2017, 2018).

### Manipulations of wrist angle and wrist angular kinematics showed relatively opposite effects on perceptual judgments of heaviness and length

To examine the effects of wrist angle and wrist angular kinematics on perception, we submitted perceptual judgments to ART ANOVA. The manipulations of wrist angle (*F*_2,1231_ = 23.56, *P* < 0.001) and kinematics (*F*_2,1231_ = 8.05, *P* < 0.001) affected perception of heaviness (Table 3). Each object was perceived to be heavier when the wrist was constrained to move about the ulnar position than the neutral and radial positions (Neutral – Ulnar, *t*_1232_ = –4.88, *η*^2^ = 0.00, *P* < 0.001; Radial – Ulnar, *t*_1232_ = –6.62, *P* < 0.001; Tables 4 & 5, and Fig. 3A, top panel), and when that object was wielded at 2Hz and 3 Hz than held statically (2 Hz dynamic – Static, *t*_1232_ = 3.96, *P* < 0.001; 3 Hz dynamic – Static, *t*_1232_ = 2.54, *P* = 0.030; Tables 4 & 5, and Fig. 3A, middle panel). Additionally, wrist angular kinematics modulated these effects of wrist angle on perception of heaviness (*F*_2,1231_ = 456.87, *P* < 0.001; Table 3). These effects of wrist angle on perception of heaviness diminished as the object was wielded at greater speeds (Radial – Ulnar: 2 Hz dynamic – Static, *P* = 0.006; Neutral– Ulnar: 3 Hz dynamic – Static, *P* = 0.007; Radial – Ulnar: 3 Hz dynamic – Static, *P* = 0.027; Tables 4 & 5, and Fig. 3A, bottom panel).

**Table 5.** Pairwise contrasts and associated Cohen’s *d*-like effect sizes following linear mixed-effects models (LMEs) that retain the model structure used in ART ANOVAs. ART ANOVAs in Table 3 are used for inference instead of LMEs as the data are not normally distributed and hence, require non-parametric tests. The outcomes of LMEs reported below provide a sense of effect sizes.

|  |  | Perceived heaviness |  |  |  | Perceived length |  |  |  |
| --- | --- | --- | --- | --- | --- | --- | --- | --- | --- |
| Effect | Contrast | Estimate ($\pm 1SE$) | Df | *t*-ratio | *d<sub>r</sub><sup>a</sup>* | Estimate ($\pm 1SE$) | Df | *t*-ratio | *d<sub>r</sub><sup>a</sup>* |
| Wrist angle |  |  |  |  |  |  |  |  |  |
|  | Neutral – Ulnar | –6.60 (3.84) | 1231 | –1.72 | –0.12 | 1.80 (0.56) | 1231 | 3.21 | 0.22 |
|  | Radial – Ulnar | –10.65 (3.84) | 1231 | –2.77 | –0.19 | 2.88 (0.56) | 1231 | 5.14 | 0.35 |
|  | Radial – Neutral | –4.04 (3.84) | 1231 | –1.05 | –0.07 | 1.08 (0.56) | 1231 | 1.93 | 0.13 |
| Wrist angular kinematics |  |  |  |  |  |  |  |  |  |
|  | 2 Hz dynamic – Static | 9.75 (3.84) | 1231 | 2.54 | 0.17 | –0.53 (0.56) | 1231 | –0.95 | –0.06 |
|  | 3 Hz dynamic – Static | 8.81 (3.84) | 1231 | 2.29 | 0.16 | 0.27 (0.56) | 1231 | 0.48 | 0.03 |
|  | 3 Hz dynamic – 2 Hz dynamic | –0.94 (3.84) | 1231 | –0.24 | –0.02 | 0.80 (0.56) | 1231 | 1.43 | 0.10 |
| Effect | Contrast | Value | Df | $\chi^2$ | *d<sub>r</sub><sup>a</sup>* | Value | Df | $\chi^2$ | *d<sub>r</sub><sup>a</sup>* |
| Wrist angle × Wrist angular kinematics |  |  |  |  |  |  |  |  |  |
|  | Neutral – Ulnar × Static – 2 Hz dynamic | 12.17 | 1 | 1.67 | 0.22 | 3.87 | 1 | 7.95 | 0.47 |
|  | Neutral – Ulnar × Static – 3 Hz dynamic | –13.86 | 1 | 2.17 | –0.25 | –1.84 | 1 | 1.79 | –0.22 |
|  | Neutral – Ulnar × 2 Hz dynamic – 3 Hz dynamic | –26.03 | 1 | 7.65 | –0.46 | –5.71 | 1 | 17.28 | –0.69 |
|  | Radial – Ulnar × Static – 2 Hz dynamic | –10.20 | 1 | 1.18 | –0.18 | –0.98 | 1 | 0.51 | –0.12 |
|  | Radial – Ulnar × Static – 3 Hz dynamic | –20.57 | 1 | 4.78 | –0.36 | –4.62 | 1 | 11.31 | –0.56 |
|  | Radial – Ulnar × 2 Hz dynamic – 3 Hz dynamic | –10.37 | 1 | 1.21 | –0.18 | –3.64 | 1 | 7.04 | –0.44 |
|  | Radial – Neutral × Static – 2 Hz dynamic | –22.37 | 1 | 5.65 | –0.40 | –4.85 | 1 | 12.46 | –0.59 |
|  | Radial – Neutral × Static – 3 Hz dynamic | –6.71 | 1 | 0.51 | –0.12 | –2.78 | 1 | 4.10 | –0.34 |
|  | Radial – Neutral × 2 Hz dynamic – 3 Hz dynamic | 15.66 | 1 | 2.77 | 0.28 | –2.07 | 1 | 2.26 | –0.25 |

<sup>a</sup>Cohen's *d*-like effect size is calculated by assuming that the sigma in classical Cohen's *d* is the residual standard deviation of the model; approach as taken in Rouder et al. (2012).

**Fig 3.**
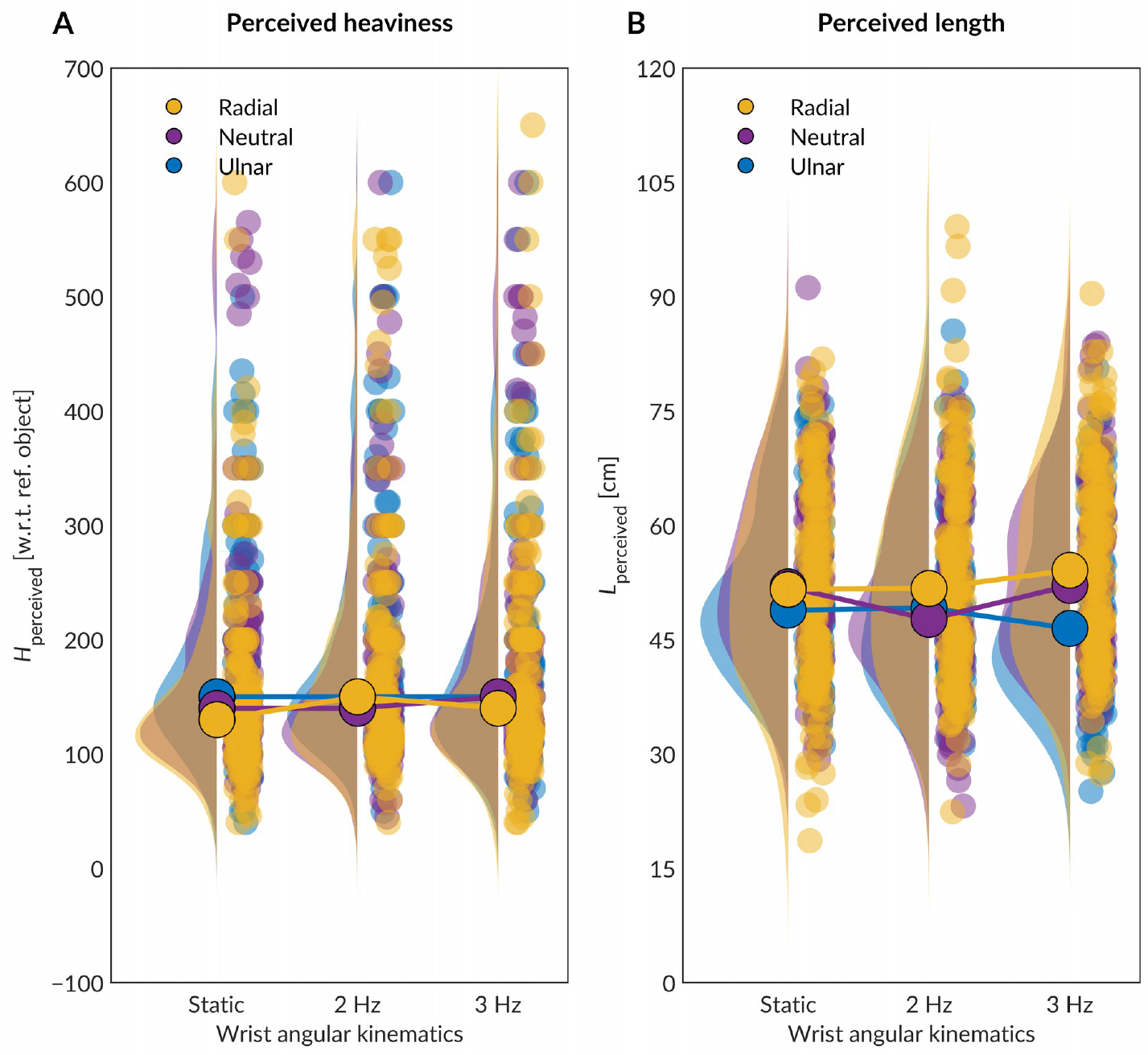
Wrist angle and wrist angular kinematics showed opposite effects on perception of heaviness and length. The solid circles describe the median value for each combination of wrist angle and wrist angular kinematics. (**A**) These effects of wrist angle on perception of heaviness diminished as the object was wielded at greater speeds (Radial – Ulnar: 2 Hz dynamic – Static, *P* = 0.006; Neutral– Ulnar: 3 Hz dynamic – Static, *P* = 0.007; Radial – Ulnar: 3 Hz dynamic – Static, *P* = 0.027). (**B**) These effects of wrist angle on perception of length amplified as the object was wielded at 2 Hz and 3 Hz than held statically (Neutral – Ulnar: 3 Hz – 2 Hz dynamic, *P* < 0.001; Radial – Ulnar: 3 Hz dynamic – Static, *P* = 0.006; Radial – Neutral: 2 Hz dynamic – Static, *P* = 0.007).

In contrast, the manipulations of wrist angle affected perception of length (*F*_2,1231_ = 13.37, *P* < 0.001), but the manipulations of wrist angular kinematics did not affect perception of length (*F*_2,1231_ = 1.36, *P* = 0.257; Table 3). Each object was perceived to be shorter when the wrist was constrained to move about the ulnar position than the neutral and radial positions (Neutral – Ulnar, *t*_1232_ = 3.55, *P* = 0.001; Radial – Ulnar, *t*_1232_ = 5.03, *P* < 0.001; Tables 4 & 5b, and Fig. 3A, middle panel). Additionally, wrist angular kinematics modulated these effects of wrist angle on perception of length (*F*_4,1231_ = 5.66, *P* < 0.001; Table 3). These effects of wrist angle on perception of length amplified as the object was wielded at 2 Hz and 3 Hz than held statically (Neutral – Ulnar: 3 Hz – 2 Hz dynamic, *P* < 0.001; Radial – Ulnar: 3 Hz dynamic – Static, *P* = 0.006; Radial – Neutral: 2 Hz dynamic – Static, *P* = 0.007; Tables 4 & 5, and Fig. 3B, bottom panel). In Table 5, we still report Cohen’s *d*-like effect sizes on the actual data (not ranked data) from the LMEs. Although the LMEs will be less reliable given the data distribution, they can still be used to make sense of the effect sizes.

In summary, manipulations of wrist angle and wrist angular kinematics affected judgments of heaviness and length in relatively opposite ways. Compared to static holding, wielding the object resulted in it being perceived heavier but wielding did not affect perceived length. As for wrist angular kinematics, wielding resulted in an object being perceived heavier but did not affect perceived length.

## Discussion

Blindfolded participants manually wielded objects of different mass and mass distribution about the wrist at different wrist angles and kinematics and reported their perceptual judgments of heaviness and length. The analysis revealed that perception of heaviness and length were predominantly dependent on an object’s static moment and the moment of inertia, respectively. Variation in judgments of heaviness and length over variation in wrist angle and wrist angular kinematics suggest that movement-related peripheral feedback from spindles and tendon organs play a fundamental role in effortful perception of both heaviness and length. However, manipulations of wrist angle and wrist angular kinematics affected perceptual judgments of heaviness and length in relatively opposite ways, suggesting that proprioceptive afferents differentially contribute to effortful perception of object heaviness and mass distribution. In what follows, we discuss possible explanations of these findings.

The present findings support Luu et al.’s (2011) unifying hypothesis in which the effortful perception of heaviness relies on both efferent and afferent information. This hypothesis takes the premise that volitional motor commands are accompanied by an expectation of reafference (i.e., sensory feedback) arising from the consequences of the motor output. Deviations from the expected reafference may be interpreted as indicating that an object is lighter or heavier. We interpret the present results related to perception of heaviness in the light of this unifying hypothesis and also reconcile several previous results using the same hypothesis.

Based on von Holst and Mittelstaedt’s reafference principle (von Holst and Mittelstaedt 1950) that proprioceptive afferents will create reafference related to the dynamics of the muscular contraction, to convey the heaviness and mass distribution of the objects to the nervous system. Briefly, afferent feedback from group Ia of muscle spindles conveys the rate of change of muscle length to the nervous system, group II muscle spindles convey a muscle’s instantaneous length (Al-Falahe et al. 1990; Proske and Gandevia 2012), and group Ib of tendon organs mainly convey the force produced by an active muscle. In our experiment, when an object was held statically, the ulnar and radial deviations of the wrist would increase tension in the muscles which produce wrist radial and ulnar deviation, as well as increase the length of the antagonists. This would primarily change the reafference from group II and Ib afferents, and as a consequence, objects were perceived heavier during ulnar deviation than during the neutral wrist angle. During the radial deviation of the wrist, it is fair to assume that input from these two afferents reversed course in all muscles (compared to ulnar deviation). As a result, objects were perceived lighter than during ulnar deviation and neutral wrist angle.

Volitional contraction of muscles while wielding an object would create strong reafference from group 1a afferents, reflecting rapid changes in muscle length. Since perceived heaviness was higher during wielding, the sensory feedback from 1a afferents may play a role in perception of heaviness. Perceived heaviness did not vary with the wrist angle when an object was wielded. This result suggests that the movement-related reafference from group 1a afferents may contribute towards perception of heaviness independent of the wrist angle (and consequently independent of input from 1b afferents). In contrast, in the absence of reafference from group Ia afferent when an object is held statically, perception of heaviness is based primarily on reafference from group Ib and group II afferents.

In the context of perception of length, the manipulations of wrist angle and wrist angular kinematics showed relatively opposite effects on perceived length, and within each pair of objects with the same static moment, the object with a greater moment of inertia was perceived to be longer. Both these findings undermine any association between a simple increase in peripheral afferent feedback and perception of length. Previous findings also indicate a lack of such association, as people can perceive the length of an object with reasonable accuracy under reduced peripheral afferent feedback: by moving an object minimally (Carello et al. 1992; Lederman et al. 1996), and by wielding an object when it is immersed in water, and the force of buoyancy reduces the force required to wield that object (Pagano and Donahue 1999; Pagano and Cabe 2003; Mangalam et al. 2017, 2018). Nonetheless, the finding that manipulations of wrist angle and wrist angular kinematics affected perceived length is important. What spatiotemporal characteristics of peripheral afferent feedback, if not simply the magnitude, may contribute to perception of length and other properties related to mass distribution?

We would also like to highlight that the two repetitions for each combination of wrist angle, wrist angular kinematics, and object are insufficient to develop a measure of central tendency for any individual. However, there are a few points to be noted here. First, we were interested in defining how individuals use proprioceptive and kinesthetic sensory feedback to estimate mass and inertial properties. We particularly wanted to avoid confounds due to long-term memory, practice and/or comparisons with previous trials. Thus, to minimize these confounds, we could use neither multiple repetitions nor long trials. Therefore, we have refrained from implying that the perceptual reports of participants imply a central tendency or a stable psychophysical measure. Second, we also want to clarify that we did not use a mean of the two values for each for each combination of wrist angle, wrist angular kinematics, and object but instead, considered both values.

Another limitation of the present study is that during the 5 s trial period, the baseline position of the wrist (either radial or ulnar) may have drifted to reach a more natural neutral position. Furthermore, the static moment and the moment of inertia of the objects may have also influenced the magnitude of drift. Although we instructed the participants to minimized the movement amplitude in the 2 Hz and 3 Hz dynamic conditions, any amount of wrist movement may have influenced the baseline wrist angle. However, the differential interaction effects of wrist angle and wrist angular kinematics on perceived heaviness and length suggest that, for the most part, this factor did not undermine the perceptual outcomes of wrist angle manipulations. The finding that we did not find any interaction effects of wrist angle, wrist angular kinematics, and object on perceived heaviness and length further supports this assertion.

Overall, the present findings suggest that effortful perception of length (or mass distribution) does depend on movement-related peripheral afferent feedback. When an object is held statically, the radial and ulnar deviations have an opposite effect on the afferent input from group II and Ib afferents of antagonist muscles. This contrast may explain why each object was perceived longer in the radial position than the neutral or ulnar positions, suggesting that during a static grasp of an object, the central processing of group II and Ib afferents together, might convey information on length (or mass distribution) to the nervous system.

It is crucial to note that the haptic perceptual system may rely on two object parameters instead of one to estimate a given object property. Relying on multiple object parameters allows deviating from a physical model, depending on the confidence in each parameter owing to the influence of exploratory conditions. For instance, when holding an object horizontally at its center of mass, perception of length can rely only on its moment of inertia. Conversely, when angular acceleration is prevented by holding an object in place vertically, perception of length has to rely on the static moment, as the moment of inertia cannot be used to perceive the length of an object that cannot be moved radially. Thus, to perceive a given object property, the haptic perceptual system may not only rely on more than one object parameter but often on different combinations of object parameters under different exploratory conditions; hence redundancy in movement-related peripheral afferent feedback is essential to flexibility and context-sensitivity of the haptic perceptual system.

In conclusion, the present findings indicate that distinct components of movement-related peripheral afferent feedback contribute to effortful perception of heaviness and mass distribution of a manually wielded object. Perception of heaviness and mass distribution is mainly derived from peripheral afferent feedback rather than central motor commands alone. An extensive investment of the central nervous system through the coupling of the fusimotor and skeletomotor control is thus fundamental to forming stable percepts of heaviness and mass distribution.

## Acknowledgments

We thank Ryan Chen and Terrence R. McHugh for help with testing the participants.

## Author contributions

M.M. and T.S. conceived and designed research; M.M. performed experiments; M.M. and N.D. analyzed data; M.M., N.D., and T.S. interpreted results of experiments; M.M. prepared figures; M.M. and T.S. drafted manuscript; M.M., N.D., and T.S. edited and revised manuscript; M.M., N.D., and T.S. approved final version of manuscript.

## Compliance with ethical standards

## Conflict of interest

The authors declare that no competing interests exist.

## Notes

### Competing Interest Statement

The authors have declared no competing interest.

